# Methylglyoxal induces nuclear accumulation of p53 and γH2AX in normal and cancer cells

**DOI:** 10.1101/2020.03.19.998773

**Authors:** Astrid Veß, Thomas Hollemann

## Abstract

The formation and accumulation of methylglyoxal (MGO) is associated with age-related diseases such as diabetes, cancer and neurodegenerative disorders. MGO is the major precursor of non-enzymatic glycation of macromolecules affecting their function and structure. We now show for the first time that MGO stress not only led to cellular aging responses like DNA double-strand breaks indicated by an accumulation of γH2AX in the nucleus. We also observed an immediate increase of Ser15 phosphorylated p53 in the nucleus of MGO treated cell lines which will change the cellular expression pattern with adverse effects on the cell cycle and other cellular functions not necessarily related to aging.

## Introduction

The formation and accumulation of methylglyoxal (MGO), the most potent glycating agent in humans, is associated with age-related diseases such as diabetes, obesity, atherosclerosis, cancer and neurodegenerative disorders [1]. Methylglyoxal is mainly formed by the non-enzymatic degradation of triose phosphates, glyceraldehyde-3-phosphate (G3P) and dihydroxyacetone-phosphate (DHAP), as a byproduct of glycolysis. It takes place in all cells and organisms. [2, 3]. The actual concentration of MGO in the cell depends on several factors like the rate of detoxification by glyoxalases, the phosphate pool of the cell and the rate of influx of carbon sources [4].

MGO is the major precursor of non-enzymatic glycation of proteins, lipids and DNA, subsequently leading to the formation of a heterogeneous group of molecules, collectively called AGEs (advanced glycation endproducts) [1, 3, 5]. The glycation reactions can cause the formation of complexes and irreversible adducts of these macromolecules affecting their function and structure. Accumulation of AGEs compromises cellular processes resulting in mitochondrial dysfunction, genomic instability, loss of proteostasis, inflammaging and cellular senescence [6]. Moreover, other studies revealed that elevated MGO levels may also have beneficial effects in cancer and lifespan [7, 8].

Tumor cells differ from non-tumor cells concerning their high glycolytic rates under anaerobic conditions (Warburg effect). This inefficient energy production, cause a high rate of glucose uptake and glycolysis in tumors resulting in elevated MGO levels [9]. Loarca *et al*. reported that treatment of hepatocellular carcinoma (HCC) cell lines with MGO promote the localization of p53 into the nucleus whereas total cellular p53 levels are not altered [10]. p53 is a well-known tumor suppressor that is mutated in many tumor cells [11]. It functions as a transcription factor, which is involved in the regulation of the cell cycle, DNA repair and apoptosis. Upon cellular stress such as DNA damage, hypoxia or viral infection p53 is activated and stabilized by different post-translational modifications interfering with its degradation [12]. It was shown that DNA damage induces phosphorylation of p53 on Ser15 [13]. This impairs the binding to MDM2, promoting the activation of p53 and thereby leading to cell cycle arrest or apoptosis [14, 15].

Using several tumor and non-tumor cell lines we showed that MGO stress led to DNA double-strand breaks, and an immediate increase of Ser15 phosphorylated p53 in the nucleus. This upregulation of nuclear phospho-p53 seems not accompanied with an increase in apoptosis rate but will likely activate p53 dependent signaling pathways and thereby change the cellular expression pattern with adverse effects on the cell cycle and other cellular functions. The application of MGO as a compound to mimic increased glycation in cells is a widely accepted method particular in the aging research field. Therefore, we suggest monitoring phosphorylated p53 and γH2AX in MGO treated cells in age related issues to exclude p53 signaling interfering with age response.

## Materials and methods

### Reagents

Methylglyoxal (MGO) was purchased from Sigma-Aldrich (catalog no. M0252). Primary antibodies used were anti-phospho-p53 (Cell Signalling, catalog no. 9286), anti-p53 (Santa Cruz), anti-phospho-Histone H2A.X (Ser139) clone JBW301 (Millipore, catalog no. 05-636), anti-cleaved caspase3 (5A1E) (Cell Signalling, catalog no.9664) and anti-PARP (Cell Signalling, catalog no.9542). F-actin was visualized with Atto 546 Phalloidin (Sigma-Aldrich). Alexa Fluor-conjugated and IRDye® (800CW, 680RD) secondary antibodies were from Thermo Fisher Scientific or Li-Cor (Bad Homburg, Germany), respectively. DNA was stained with DAPI (Sigma-Aldrich).

### Cell Culture and Methylglyoxal (MGO) treatment

The breast cancer cell line MDA-MB-468 was cultured at 37°C and 5% CO2 in RPMI medium supplemented with 10% (v/v) fetal calf serum (FCS) and Antibiotic-Antimycotic (Thermo Fisher Scientific, Schwerte, Germany). The keratinocyte cell line HaCaT, cervical adenocarcinoma Hela cells and neuroblastoma SH-SY5Y cells were cultured at 37°C and 5% CO2 in DMEM high glucose medium supplemented with 10% (v/v) fetal calf serum (FCS), 10% sodium pyruvate and Antibiotic-Antimycotic (Thermo Fisher Scientific). Primary human fibroblasts were cultured at 37°C and 5% CO2 in Dulbecco’s Modified Eagle Medium (DMEM)/Hams F12 supplemented with 8% (v/v) fetal calf serum (FCS), 2% Ultroser G (PALL Life Sciences) and 2 mM L-glutamine (Thermo Fisher Scientific).

In all experiments Methylglyoxal (MGO) (Sigma-Aldrich) was diluted to 0.1 mM, 0.5 mM or 1 mM and the cells were incubated for 1h and 18h respectively.

### Immunofluorescence and Microscopy

For immunofluorescence analysis, 8×10^4^ cells (12-well plates) were grown on glass coverslips, treated with 0.5 mM or 1 mM MGO for 1h and 18h respectively, washed twice with PBS, fixed with 3.7% formaldehyde in PBS for 15 min, permeabilized with 0.2% (v/v) Triton X-100 in PBS for 10 min and blocked in 10% FCS/1% BSA/0.05% Triton X-100 (v/v) in PBS for 30 min. Primary antibodies were diluted 1:200 and incubated for 1 h at room temperature. Alexa-conjugated secondary antibodies were diluted 1:200 and applied for 1 h at room temperature. Samples were covered with ProLong Gold antifade reagent (Life Technologies) and imaged using an Apotome-containing Axio Observer.Z1 (Zeiss, Jena, Germany) equipped with a 63x oil objective and a monochrome Axiocam MRm camera. Representative images are shown.

### Apoptosis assay

2×10^5^ cells were subjected to the Caspase-3/CPP32 Fluorometric Assay Kit (BioVision) according to the manufacturer’s protocol. Absorbance at 505 nm was measured after 2 h of incubation at 37°C using a microplate reader. Quantitative results were calculated from at least three independent biological experiments.

### Western Blot

Cells were washed twice with ice-cold phosphate-buffered saline and lysed in RIPA buffer (50 mM Tris-HCl, pH 7.4, 150 mM NaCl, 2 mM EDTA, 1% (v/v) Triton X-100, 0.1% SDS, Complete EDTA-free protease inhibitor cocktail) for 30 min on ice. The proteins were separated on SDS-PAGE, transferred to polyvinylidene fluoride (PVDF) membranes (Merck Millipore), incubated with indicated primary antibodies used at 1:200–1:5000 dilutions and incubated overnight at 4°C. Fluorophore-labeled secondary antibodies used at 1:10.000 dilutions (LI-COR, Bad Homburg, Germany) were incubated for 1 h at room temperature. The fluorescence signals were detected with ODYSSEY CLx (LI-COR) and quantified by the associated software. Quantitative results were calculated from at least three independent biological experiments.

## Results and Discussion

In this study we analyzed the impact of the glycating agent Methylglyoxal on the phosphorylation of p53 at Ser-15 and its subsequent translocation to the nucleus. We compared different non-tumor and tumor cell lines to figure out if this response to MGO is rather a more general than a cell-type specific (depending on the genetic background) answer. Here we treated non-tumor cell lines (HaCaT keratinocytes, primary fibroblasts) and tumor cells (cervical adenocarcinoma Hela cells, breast cancer MDA-MB-468 cells and neuroblastoma SH-SY5Y cells) with concentrations of MGO between 0.1 mM and 1 mM which are generally used in this research field. The localization of Ser-15 phosphorylated p53 was analyzed 1 h and 18 h after treatment in the above mentioned cell lines. In non-tumor cells (HaCaT cells) as well as in tumor cells (MDA-MB-468 cells) MGO treatment caused a significant increase of Ser15 phosphorylated p53 in the nucleus (Fig 1A-D). We found that MGO increased nuclear phospho-p53 levels up to ∼67 fold in HaCaT cells and up to ∼1.8 fold in MDA-MB-468 cells (Fig 2). However, in Hela cells and primary human fibroblasts, MGO treatment had no effect on the localization of phospho-p53 (S1 Fig; data not shown) and in SH-SY5Y cells the nuclear localization of phospho-p53 is already very high without MGO treatment (data not shown).

**Fig 1.**
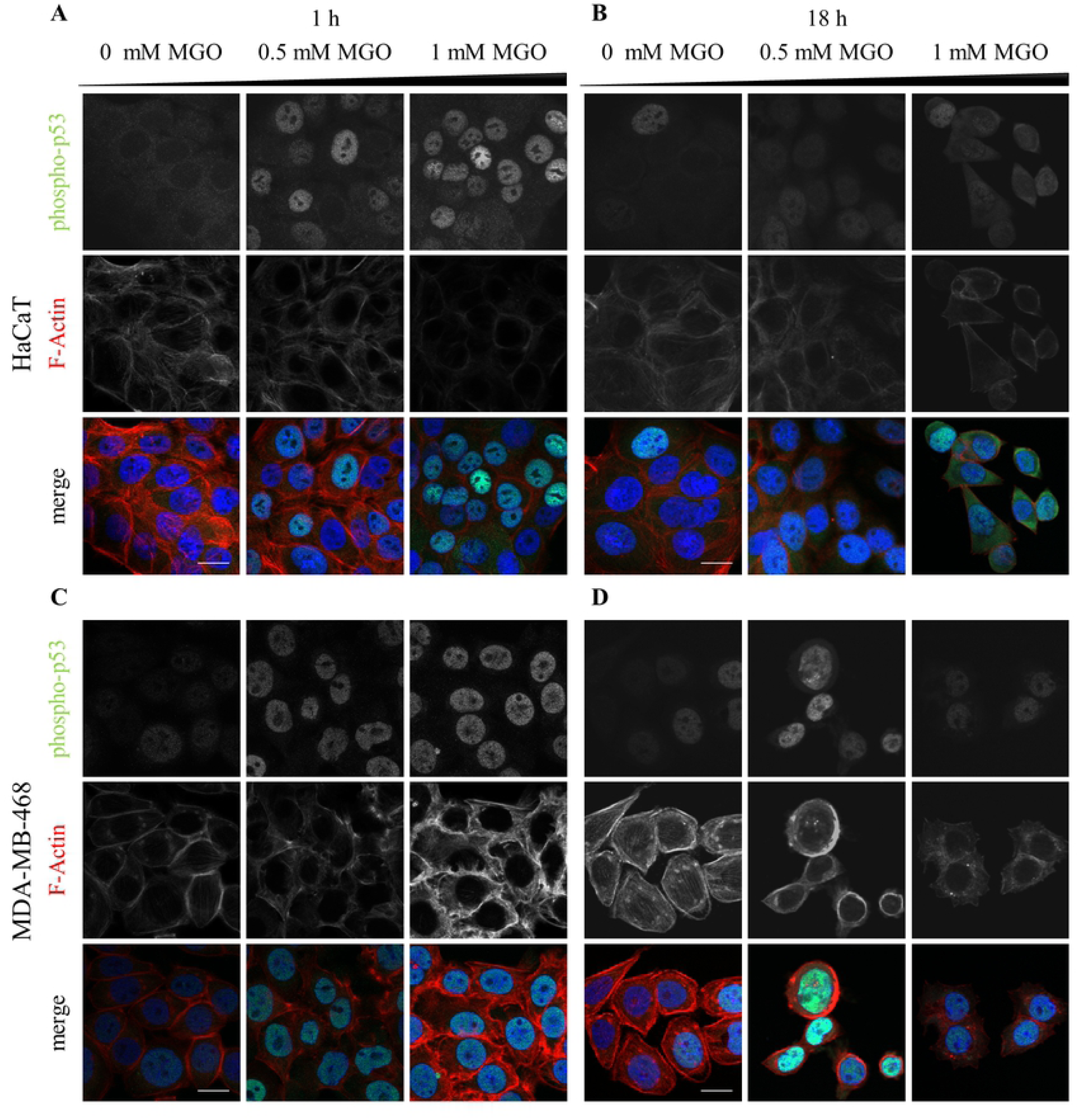
Analysis of the expression and localization of phospho-p53 in HaCaT cells and breast cancer MDA-MB-468 cells after treatment with MGO. Representative images of HaCaT (A,B), MDA-MB-468 cells (C,D) treated with indicated MGO concentrations for 1h (A,C) and 18 h (B,D). The cells were stained for phospho-p53 together with phalloidin (F-actin). Merged images show phospho-p53 (green) and F-actin (red). Scale bars, 20 µm.

**Fig 2.**
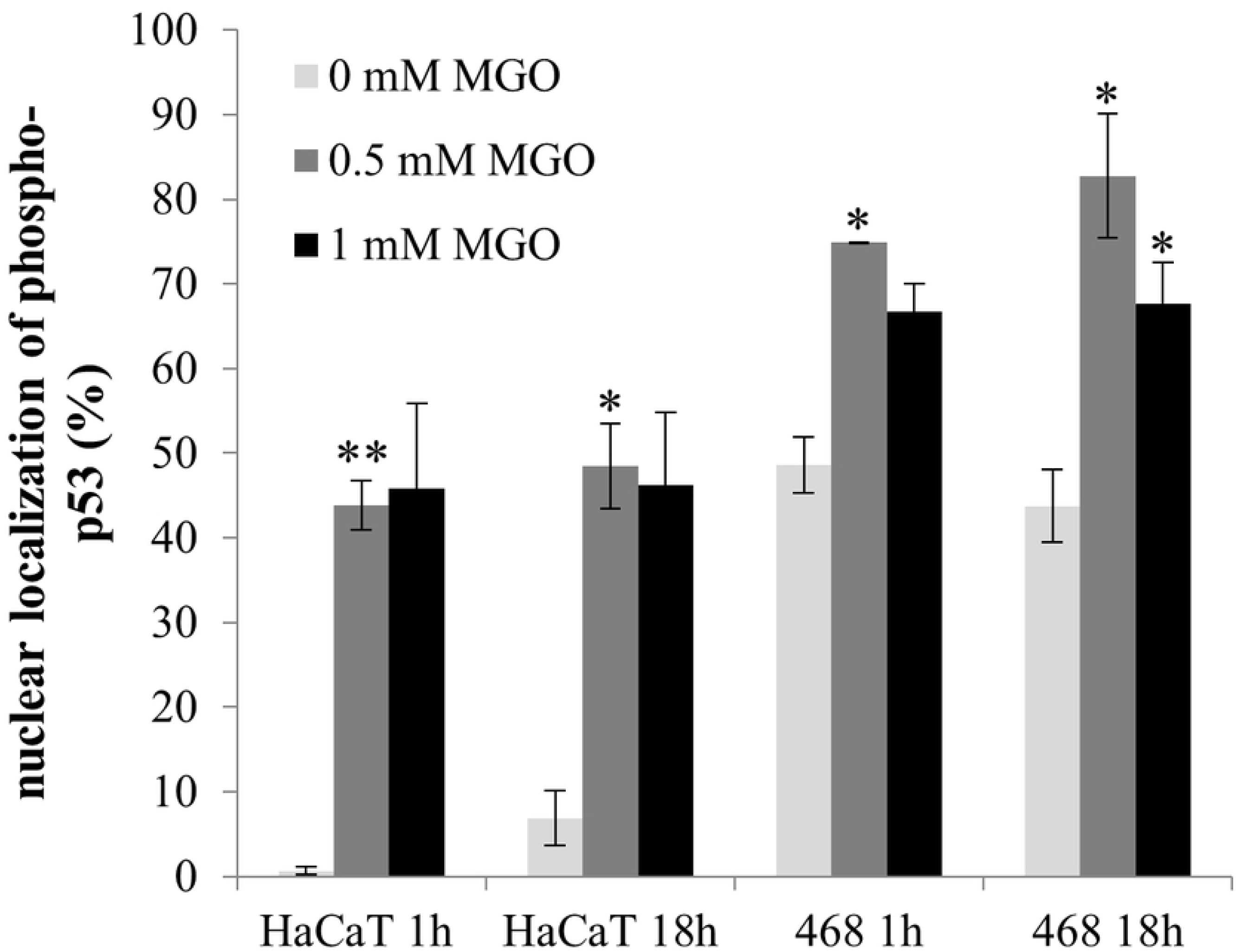
MGO treatment results in an increase in nuclear localized phospho-p53 in HaCaT and MDA-MB-468 cells. Percentage of cells showing nuclear localization of phospho-p53 normalized to the total cell number. All error bars represent ± SEM, n≥2, N>50. All *P*-values were calculated using an unpaired two-sample Student’s *t*-test (\**P*≤0.05, \*\**P*≤0.01).

It was shown previously that MGO is a cytotoxic agent, which can lead to apoptosis in cancer cells through the generation of ROS [16], DNA modification and DNA-protein crosslink [17]. To answer the question if DNA damage esp. DSBs induced by MGO is a prerequisite for the increase in nuclear localization of phospho-p53 we determined the phosphorylation level of histone H2AX in HaCaT cells and human primary fibroblasts upon MGO treatment. Phosphorylated H2AX (γH2AX) specifically binds to damaged DNA and can be visualized via immunofluorescence staining. Upon binding of γH2AX to damaged DNA several proteins are recruited for instance BRCA1 and p53 binding protein1 to induce DNA repair [18]. We could demonstrate that MGO treatment already after 1 h increased nuclear γH2AX accumulation in HaCaT cells and human primary fibroblasts (Fig 3 and 4). Since in human primary fibroblasts, MGO treatment does not lead to nuclear p53 translocation, DNA double strand breaks seem to be not sufficient to trigger nuclear localization of phospho-p53.

**Fig 3.**
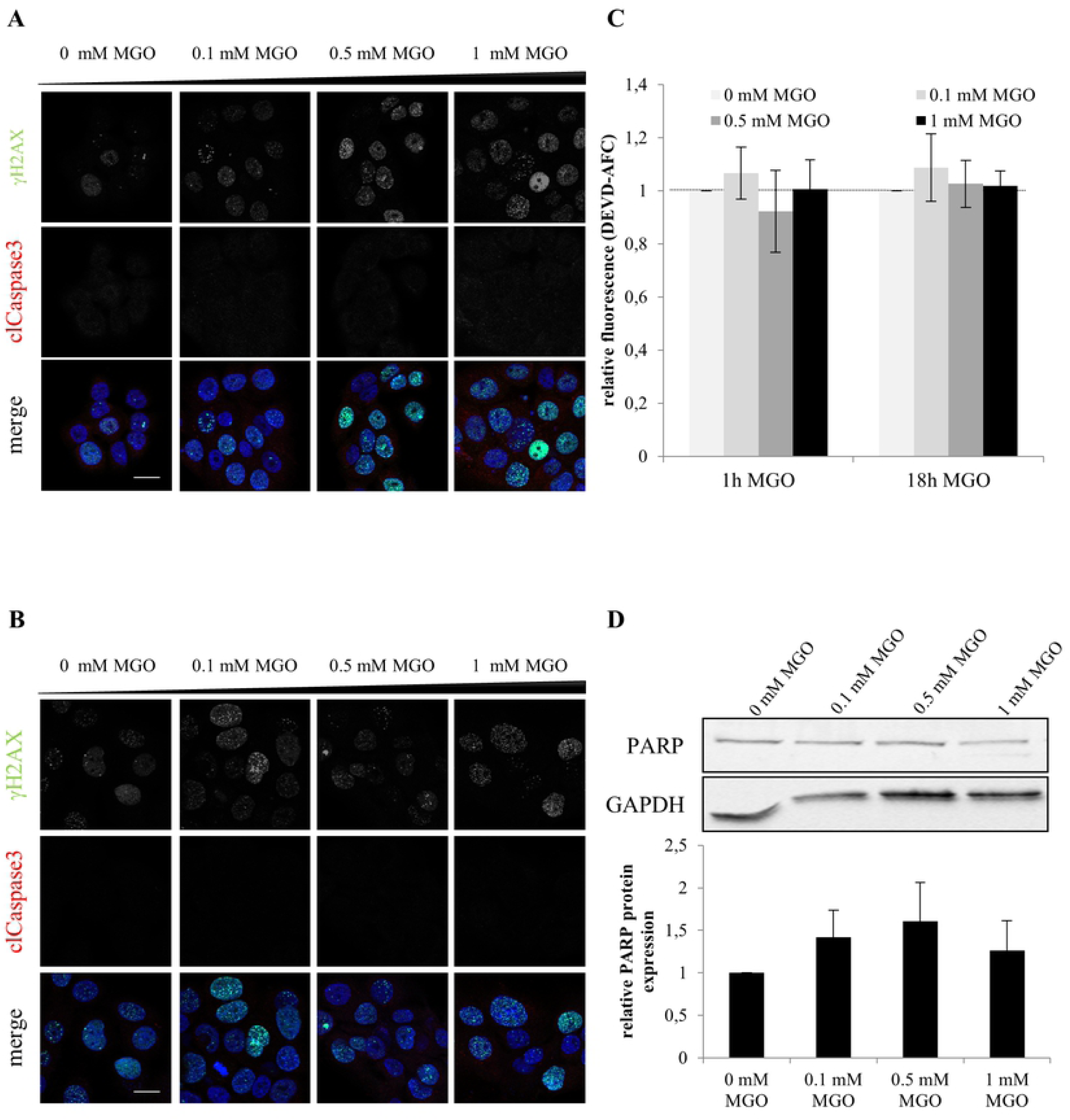
Analysis of DNA double-strand breaks and apoptosis rate in HaCaT cells after treatment with MGO. Representative images of HaCaT cells treated with indicated MGO concentrations for 1h (A) and 18h (B). The cells were stained for γH2AX together with cleaved caspase. Merged images show γH2AX (green), cleaved caspase (red) and DNA (blue). Scale bars, 20 µm. C. Apoptosis rate in HaCaT cells by measuring caspase-3 activity using the Caspase-3/CPP32 Fluorometric Assay Kit. All error bars represent ± SEM, n=3. D. Representative western blot of PARP in total cell lysates treated with indicated MGO concentrations for 18h. GAPDH was used as a loading control. Quantification of PARP protein levels normalized to GAPDH. All error bars represent ± SEM, n=3.

**Fig 4.**
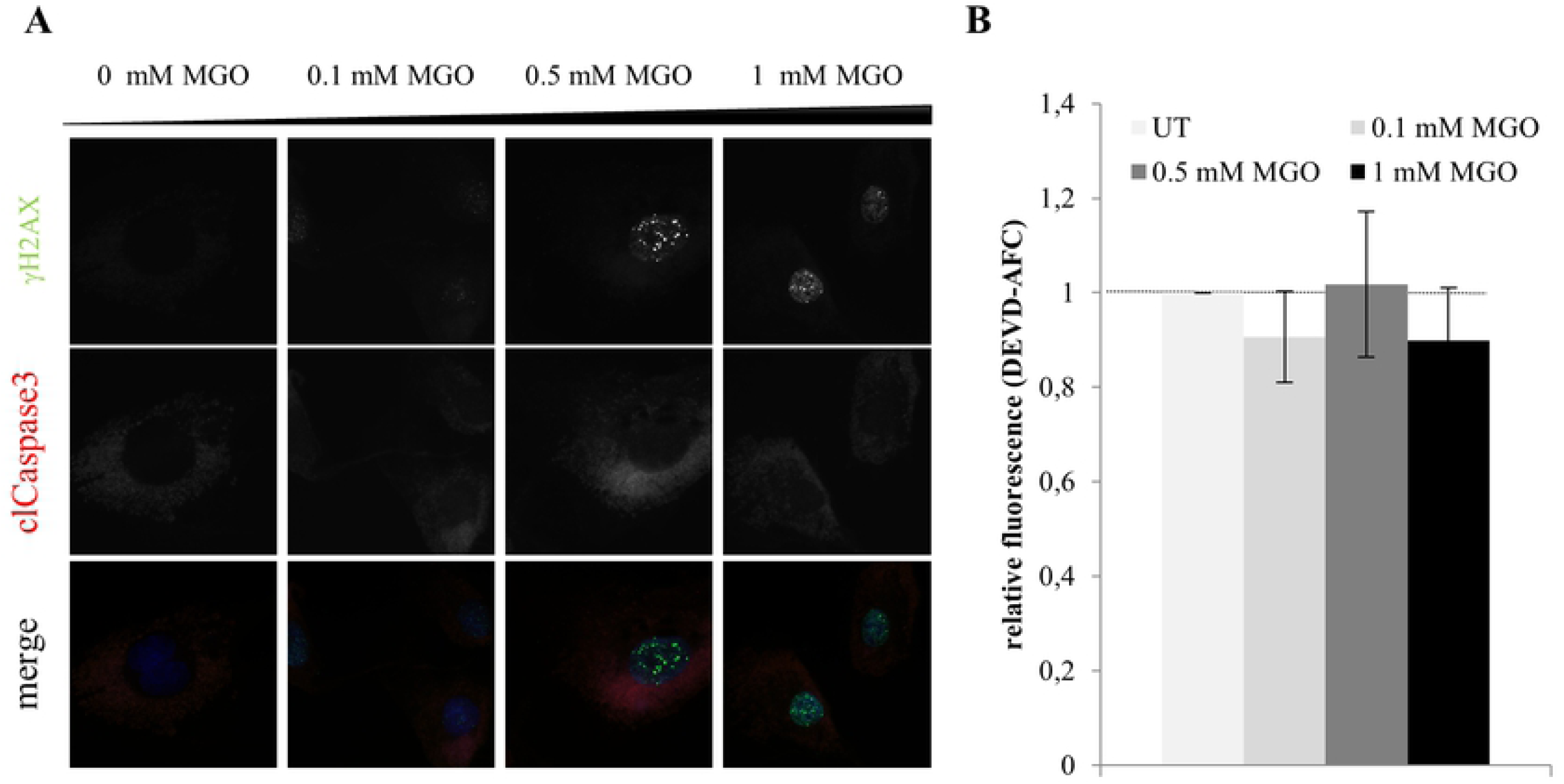
Analysis of DNA double-strand breaks and apoptosis rate in primary human fibroblasts after treatment with MGO. A. Representative images of primary human fibroblasts treated with indicated MGO concentrations for 1h. The cells were stained for γH2AX together with cleaved caspase. Merged images show γH2AX (green), cleaved caspase (red) and DNA (blue). Scale bars, 20 µm. B. Apoptosis rate in primary human fibroblasts after treatment with indicated MGO concentrations for 1 h by measuring caspase-3 activity using the Caspase-3/CPP32 Fluorometric Assay Kit. All error bars represent ± SEM, n=3.

High levels of activated p53 may drive cells into an apoptotic fate. Thus we asked whether the number of apoptotic cells is altered upon MGO treatment. To this end, expression and activity of Caspase3 and PARP cleavage was investigated. We could demonstrate that upon treatment of HaCaT cells and human primary fibroblasts with MGO the apoptotic rate was not significantly changed (Fig 3A-C, Fig 4). Also PARP as one of the main cleavage target of caspase3 in vivo ([19]) was not affected (Fig 3D).

p53 is activated upon many different stress factors influencing important cellular processes either positively or negatively depending on the cellular context [20, 21]. Therefore, p53 is thought be required for the maintenance of cellular and particularly genomic integrity. This is supported by the finding, that 50% of human tumors carry loss of function mutations in the p53 gene [11, 22-25]. 95% of these detected mutations are located in the DNA binding domain of p53 [26]. In the cell lines we used in this study MGO treatment leads to different p53 responses. Only in HaCat and MDA-MB-468 cells MGO treatment causes increased accumulation of Ser15 phosphorylated p53 in the nucleus. Interestingly in these cell lines the p53 gene carries mutations in the DNA binding domain altering its ability to bind to p53 response elements and thereby changing its transcriptional activity. The MDA-MB-468 cell line is hemizygous for a mutated p53 gene (R273C) [11, 27]. Whereas the spontaneously immortalized skin keratinocyte cell line, HaCaT, possesses three point mutations (H179Y, D281G, R282W) of p53 on both alleles [28]. In contrast to that, Hela cells and probably also primary fibroblasts which show no change in the nuclear localization of Ser15 phosphorylated p53 possess exclusively wild type allels of p53. In these cells it is supposed that other mechanisms may lead to a carcinogenic inactivation of p53, for instance HPV infection for cervical carcinoma or increase in protein stability in neuroblastoma [29, 30].

In summary, in this study we showed for the first time that MGO stress leads to an immediate increase of Ser15 phosphorylated p53 in the nucleus of several cell lines. This upregulation of nuclear phospho-p53 seems not accompanied with an increase in apoptosis rate but will likely activate p53 dependent signaling pathways. This would fit with recently published data showing an activation of the p53 pathway upon MGO treatment in human umbilical vein endothelial cells (HUVECs) [31]. Moreover, it was shown that the total amount of nuclear p53 is increased in hepatocellular carcinoma cells upon MGO treatment. However, the type of posttranslational modification of p53 was not analyzed. The authors correlate the nuclear increase of p53 upon MGO treatment with a decreased migration, invasion and adhesion phenotype in these cells [10].

The application of MGO as a compound to mimic increased glycation in cells is a widely accepted method particular in the aging research field. Nevertheless, besides the enormous progress in understanding AGE related phenotypes and diseases in more detail some critical aspects of MGO treatment arises. It is important that the used concentrations of MGO are comparable to the actual physiological concentration in this specific tissue and blood respectively [1]. By using UPLC-MS/MS plasma levels of MGO in healthy individuals have been estimated at ∼60-250 nM, cellular levels ranges from ∼1-5 µM MGO [1, 32, 33]. In cell culture model systems often near toxic concentrations of MGO ranging from 50 µM up to 2 mM MGO are applied. For instance, HUVEC cells were treated with up to 800 µM MGO to copy injuries in endothelial cells typically seen in diabetes patients knowing that these concentrations do not reflect the plasma concentrations of MGO even in diabetes patients [31]. The authors argued that the in vivo situation is rather a complex mixture of compounds acting in synergy than a single compound leading to this phenotype [31]. In addition, the application of MGO in cell culture is different compared to the in vivo situation and thereby MGO can act more direct on the cells. To exclude unphysiological responses of MGO treatment in general, we suggest establishing a protocol for each cell line utilized, which includes the analysis of the Ser-15 phosphorylated p53 and γH2AX status to exclude undesirably effects of MGO which would otherwise overlap with aging mechanisms (Fig 5).

**Fig 5.**
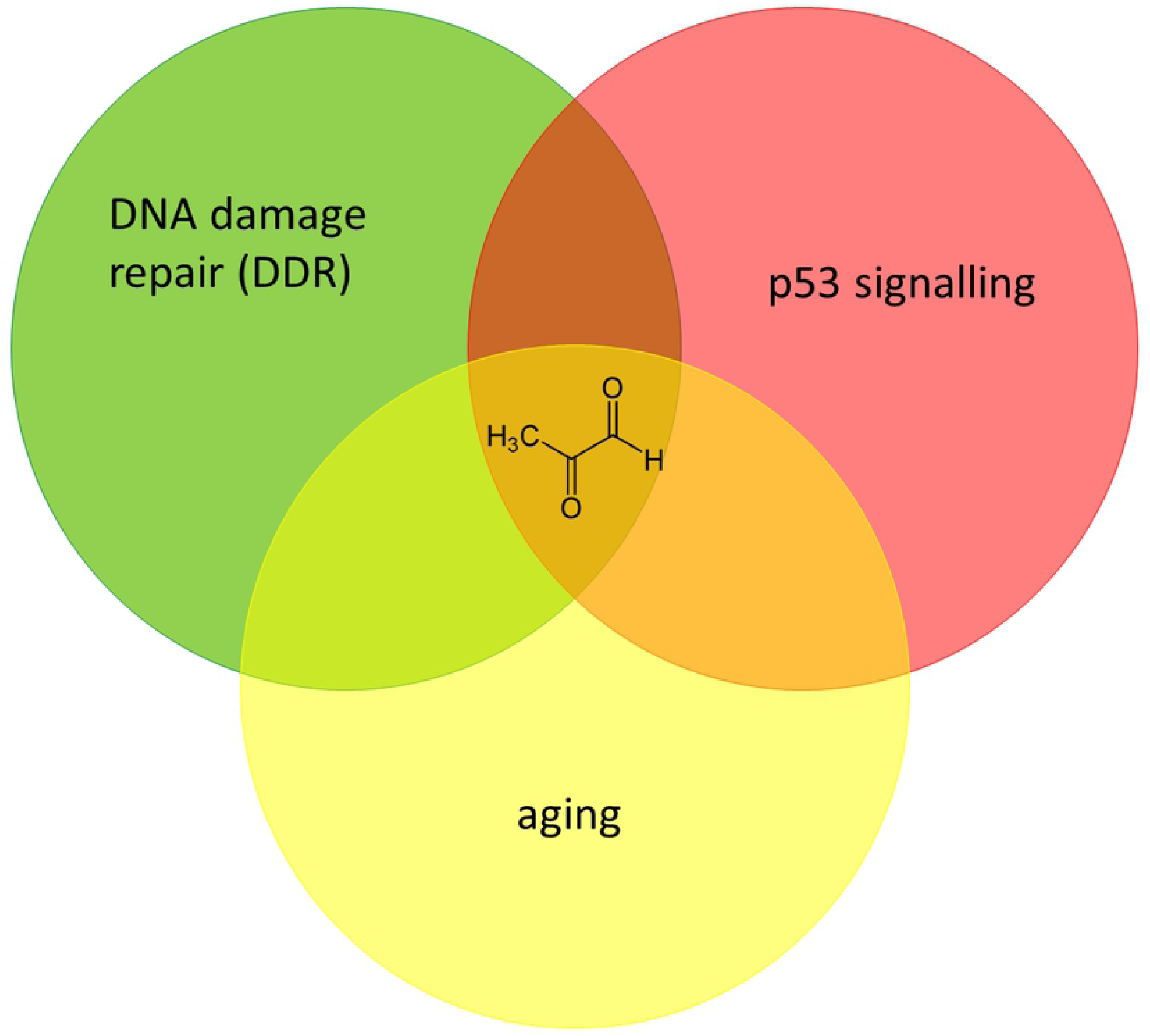
Model of MGO induced stress response. MGO stress leads to DNA double-strand breaks, and an immediate increase of phosphorylated p53 in the nucleus of MGO treated cell lines. The subsequent activation of p53 dependent signaling pathways is supposed to overlap with the aging response in these cells. We suggest monitoring phosphorylated p53 and γH2AX in MGO treated cells to exclude undesirable effects of MGO interfering with aging response.

## Acknowledgement

We thank Dr. Herbert Neuhaus for critical reading the manuscript.

## Supporting information

**S1 Fig. Analysis of the expression and localization of phospho-p53 in primary human fibroblasts after treatment with MGO.** Representative images of primary human fibroblasts treated with indicated MGO concentrations for 1h (A) and 18 h (B). The cells were stained for phospho-p53 together with phalloidin (F-actin). Merged images show phospho-p53 (green) and F-actin (red). Scale bars, 20 µm.

